# Using spatial genetics to quantify mosquito dispersal for control programs

**DOI:** 10.1101/2020.03.30.017301

**Authors:** I Filipović, HC Hapuarachchi, WP Tien, ABAR Muhammed, C Lee, CH Tan, GJ Devine, G Rašić

**Affiliations:** Mosquito Control Laboratory, QIMR Berghofer Medical Research Institute, 300 Herston Road, 4006 Herston, QLD, Australia; Environmental Health Institute, National Environment Agency, 11, Biopolis Way, #06-05-08, Singapore, 138667, Singapore

## Abstract

**Background:** Hundreds of millions of people get a mosquito-borne disease every year, of which nearly one million die. Mosquito-borne diseases are primarily controlled and mitigated through the control of mosquito vectors. Accurately quantified mosquito dispersal in a given landscape is critical for the design and optimization of the control programs, yet the field experiments that measure dispersal of mosquitoes recaptured at certain distances from the release point (mark-release-recapture MRR studies) are challenging for such small insects and often unrepresentative of the insect’s true field behavior. Using Singapore as a study site, we show how mosquito dispersal patterns can be characterized from the spatial analyses of genetic relatedness among individuals sampled over a short time span without interruption of their natural behaviors.

**Methods and Findings:** We captured ovipositing females of *Aedes aegypti*, a major arboviral disease vector, across floors of high-rise apartment blocks and genotyped them using thousands of genome-wide SNP markers. We developed a methodology that produces a dispersal kernel for distance that results from one generation of successful breeding (effective dispersal), using the distances separating full siblings, 2^nd^ and 3^rd^ degree relatives (close kin). In Singapore, the estimated dispersal distance kernel was exponential (Laplacian), giving the mean effective dispersal distance (and dispersal kernel spread σ) of 45.2 m (95%CI: 39.7-51.3 m), and 10% probability of dispersal >100 m (95%CI: 92-117 m). Our genetic-based estimates matched the parametrized dispersal kernels from the previously reported MRR experiments. If few close-kin are captured, a conventional genetic isolation-by-distance analysis can be used, and we show that it can produce σ estimates congruent with the close-kin method, conditioned on the accurate estimation of effective population density. We also show that genetic patch size, estimated with the spatial autocorrelation analysis, reflects the spatial extent of the dispersal kernel ‘tail’ that influences e.g. predictions of critical radii of release zones and *Wolbachia* wave speed in mosquito replacement programs.

**Conclusions:** We demonstrate that spatial genetics (the newly developed close-kin analysis, and conventional IBD and spatial autocorrelation analyses) can provide a detailed and robust characterization of mosquito dispersal that can guide operational vector control decisions. With the decreasing cost of next generation sequencing, acquisition of spatial genetic data will become increasingly accessible, and given the complexities and criticisms of conventional MRR methods, but the central role of dispersal measures in vector control programs, we recommend genetic-based dispersal characterization as the more desirable means of parameterization.

## Introduction

Mosquitoes’ ability to carry and transmit human pathogens (malaria and filarial parasites, arboviruses) causes hundreds of millions of infections and nearly one million fatalities every year [1]. Both prevention and mitigation of many mosquito-borne disease outbreaks are primarily reliant on the control of mosquito vectors. Most of these interventions are designed to impact mosquito abundance or daily survival by targeting immature and adult stages. In the case of the major arbovirus vector *Aedes aegypti*, an urban-dwelling container-breeding mosquito, conventional control approaches include removal and treatment of larval habitats, as well as elimination of adults through insecticide application (indoor-residual and space spraying) and lethal trapping [2]. New biocontrol strategies undergoing field evaluations include RIDL^R^ and *Wolbachia-*based population replacement and suppression [3–5].

Defining the optimal area for treatment or mosquito release is one of the key considerations when implementing a public health intervention or designing a field-trial for a new control approach. For example, to contain the spread of dengue virus during an outbreak, focal insecticide-based control of *Ae. aegypti* adults is typically conducted at and around the main and secondary residences of dengue cases. The radius of the area to be treated is informed by the average dispersal distance of potentially infected female *Ae. aegypti* [6]. Understanding the ability of released sterile male mosquitoes to disperse and mate in an area being targeted by a suppression strategy is essential for predicting the required release pattern [7]. Additionally, sustained suppression in a target zone is difficult if a surrounding buffer zone is too small to prevent immigration by gravid wild-type females from neighboring areas. Similarly, stable introduction of a virus-blocking *Wolbachia* may fail if the release area of *Wolbachia-*infected *Ae. aegypti* is too small and too vulnerable to immigration by wild-type mosquitoes [8]. For the emerging genetic-based control approaches such as gene drive systems [9], well characterized mosquito dispersal is crucial for addressing the biosafety concerns around the systems’ confineability and reversibility in the field [10,11].

Quantifying mosquito dispersal of both wild type and introduced mosquito strains in any given landscape is, therefore, critical for the considerations of the size of the treatment area and the surrounding buffer zones. Those considerations complement practical operational deliberations of the availability of human and economic resources for implementation and the sample sizes required to capture epidemiological endpoints [12,13].

Mosquito dispersal characteristics have been typically studied using conventional mark-release-recapture (MRR) experiments utilizing powders and paints on trapped or laboratory-reared adult mosquitoes [14]. Location of the recaptured marked insects relative to their release point is typically used to estimate the mean distance travelled (MDT), and the distance within which 50% or 90% of mosquitoes are expected to disperse (FR50 and FR90, respectively). Fewer MRR studies have incorporated the dispersal kernel theory to estimate the distribution of dispersal distances over the whole flight range [7,15,16]. MRR experiments in *Ae. aegypti* have reported the mean dispersal distance to range from tens to hundreds of meters [14], and this variation points to a need to characterize dispersal locally so that the optimal control can be deployed in a given landscape. However, MRR experiments are operationally demanding, and the rearing and marking procedure can alter mosquito fitness and movement behavior in the field [17]. Additionally, the release of biting vector-competent females might pose an unacceptable risk of increased pathogen transmission in endemic areas.

Here we show how information on mosquito dispersal characteristics can be obtained from the spatial patterns of genetic distance and relatedness among sampled individuals, providing an alternative to the MRR experiments for informing the mosquito control programs. In contrast to conventional MRR approaches that require insect trapping or rearing, followed by mark, release and recapture, the genetic approach requires only insect capture, utilising the information from genetic markers and spatial location of individuals sampled continuously across an area over a limited time span and without manipulation or interruption of their natural behaviors.

In social insects like bumblebees, queen dispersal distance has been estimated by comparing the locations of workers (sampled in summer) and queens (sampled in the following spring) that were identified as sisters through sibling reconstruction analysis with genetic markers [18]. Inferences about mosquito ovipositing behavior have also been made using the genetic reconstruction of sibling groups, where the distance between full siblings sampled from different larval sites directly reflects mother’s movement distance during her skip-oviposition [19–21]. Here we show that the distance separating not only full siblings (1^st^ degree relatives), but also 2^nd^ and 3^rd^ degree relatives (close-kin), can be used to estimate the dispersal distance over one generation of successful breeding (i.e. effective dispersal distance) in insects like *Ae. aegypti*.

Our newly developed method decomposes the observed separation distances between close-kin (sampled as breeding adults) to generate the distribution of potential effective dispersal distances and to parametrize the dispersal distance kernel. This dispersal kernel provides the density of probability that a dispersal event ends at a given distance away from the source, regardless of the direction. Importantly, it refers to the dispersal distance achieved over one generation of successful reproduction, such as distance between the birthplace and the ovipositing place of a female (i.e. distance between the ovipositing place of a mother and a daughter). We demonstrate the robustness of the method to produce dispersal kernel parameters consistent among different subsets of data (with one or multiple kinship categories) and congruent with the estimates of dispersal characteristics from the previously published MRR experiments in *Ae. aegypti*.

When few close kin are captured, the conventional genetic analysis of isolation-by-distance (IBD) between unrelated individuals can be used to estimate the spread of the dispersal kernel from the slope of the IBD relationship and the effective density, and we show that its results can be congruent with the new close-kin method. Finally, we show that the estimated genetic patch size from the spatial autocorrelation analysis reflects the spatial extent of the effective dispersal distance kernel’s ‘tail’ that cannot be ascertained with IBD analysis alone.

We analyzed the genotyped and geo-referenced *Aedes aegypti* individuals collected in two densely populated areas of Singapore with a homogeneous distribution of high-rise apartment blocks. *Aedes aegypti* is the primary vector of dengue virus in Singapore that, despite having a low *Aedes* house index (2%) and an extensive vector surveillance and control program [22], continues to experience regular dengue outbreaks [23,24]. This dataset offered an opportunity to characterize *Ae. aegypti* dispersal in a highly urbanized landscape with a prominent vertical dimension, but the spatio-genetic analytical approach (the new close-kin method, and the conventional IBD and spatial autocorrelation analyses) can be applied across different landscapes and vector species.

## Materials and Methods

### Field collections

Adult *Aedes aegypti* females were collected using the sticky traps developed by the Environmental Health Institute, National Environment Agency of Singapore (EHI, NEA), known as the Gravitraps - simple, hay infusion-filled cylindrical traps with a sticky inner surface that preferentially catch gravid females in search of suitable ovipositing sites [25]. The Gravitraps have been deployed in public housing estates island-wide since 2017 as part of the vector surveillance program. For the current study, we chose two public housing estates with high-rise blocks: Tampines (30 acre sampling area in patch 1, 10 acres in patch 2) and Yishun (46 acre sampling area) that are 14 km apart (Suppl. Fig. 1), with the Gravitraps positioned for vertical sampling in each building: ground level (1^st^-2^nd^ floor), mid-level (4^th^-5^th^ floor), and high level (9^th^ floor and above). All analysed individuals were collected in weekly intervals between April 1 and May 30 2018 and their geo-locations and vertical positions were recorded.

### DNA extraction. sequencing and genotyping of individual mosquitoes

Total genomic DNA was extracted from individual mosquitoes using Dneasy Blood and Tissue DNA extraction kit (Qiagen, Hilden, Germany), quantified with the Qubit High Sensitivity DNA kit (Thermo Fisher Scientific, Waltham, MA, USA) and 100 ng was used for downstream processing. Reduced-genome representation sequence data were generated for each individual using the double-digest RAD sequencing approach by Peterson et al. [26], with the sample processing and library preparation protocol as described in Rašić et al. [27]). ddRAD-seq libraries were sequenced on the Illumina HiSeq4000 platform. The sequencing data were processed (trimmed to 90 bp and filtered for quality) using the bash script/pipeline from Rašić et al. [27], and the high-quality reads were aligned to the L5 version of the *Ae. aegypti* genome assembly [28] using Bowtie [29]. Unambiguously mapped reads converted to bam format were processed in SAMtools [30] to generate sorted bam files that were used to produce genotype likelihood files and VCF using the SAMtools variant calling method as implemented in ANGSD [31]. The final dataset included 107 mosquitoes from Tampines and 108 from Yishun, that had <30% missing data at 83,255 and 69,051 variable sites (SNPs) for the Tampines and Yishun datasets, respectively.

### Inference of familial relationship (kinship estimation)

Relationships between individuals were determined using the recently-developed approach by Waples et al.[32] as implemented in the program NGSRelate [33]. This method shows improved accuracy and precision when compared to related approaches, given that (a) it does not require population allele frequency estimates - instead, the framework calculates two-dimensional site-frequency-spectrum for each pair of individuals, and (b) it is applied directly to sequencing data (*via* genotype likelihoods) rather than the called genotypes, which is particularly suitable for lower-depth sequencing data acquired in RAD-seq experiments [32]. Between 40K and 48K SNPs were used by NGSRelate in each of the pairwise calculations across the entire dataset. For the spatial analyses, we considered close kin as pairs with an inferred category of 1^st^, 2^nd^ or 3^rd^ degree relatives. Kinship categories were determined based on the combination of three statistics (R0, R1 and KING-robust kinship), that allows the distinction between the parent-offspring and the full-sibling relationship within the category of 1^st^ degree relatives, which is difficult to achieve with other available methods [32]. Second degree relatives include half-siblings, avuncular and grandparent-grandoffspring pairs that cannot be genetically distinguished, but in our sampling scheme (collection of gravid females over a short time period) the grandparent-grandoffspring category is the least likely. Also, we can assume that half-siblings are paternal (i.e. half-sisters share a father, not a mother) given that *Ae. aegypti* females rarely mate more than once [34,35]. We assume that 3^rd^ degree relatives are first cousins, given that a category like great-grandparent/great-grandoffspring is very unlikely in our sampling scheme.[32]

### Genetic and geographic distance among individuals

We used different individual-based genetic distances among individuals within each area (Yishun or Tampines). The PCA-genetic distance was estimated by first performing the Principal Component Analysis (PCA) from genotype data in the R package ‘adegenet’ [36] and then creating a distance matrix from the Euclidean distance among the maximal number of PC axes. PCA genetic distance does not assume any particular microevolutionary processes and it exhibits a linear relationship with Euclidean geographic distance, showing the highest model selection accuracy in landscape genetic studies, especially when dispersal rates are high across the examined area [37]. We also estimated Rousset’s genetic distance *â* [38] and Loiselle’s kinship coefficient [39] in the program SPAGEDI [40].

Pairwise spatial (geographic) distance between mosquitoes was calculated as the shortest straight line (Euclidean) distance in three dimensions, based on the X/Y (long/lat) and Z (height) coordinates of their collection point, here called the Euclidean 3D distance, represents a linear geographic distance in meters (m). Natural logarithm (ln) of this distance was used in the analyses where Rousset’s *â* or Loiselle’s kinship coefficient was applied (see below), given that both of these genetic coefficients exhibit approximate linear relationship with ln-geographic distance [40].

### Estimation of mosquito dispersal characteristics

Dispersal characteristics of *Ae. aegypti* were estimated in three ways: (1) by constructing a dispersal kernel based on the recorded distances between close kin (CK), (2) using the isolation-by-distance (IBD) framework, and (3) spatial autocorrelation analysis.

### Dispersal kernel estimation from close-kin data

We parametrized the dispersal kernel using the estimates of the effective dispersal distance that we define as a distance between the birthplace and the ovipositing place of a female.

In our sampling scheme, adult females were caught after landing on a lethal ovipositing site (Gravitrap) and we assume that this is a result of their active flight (and not passive, human-assisted movement). We consider pairs of females caught in different traps that could be genetically assigned to one of the following kinship categories: parent-offspring (*po*), full siblings (*fs*), 2^nd^ degree relatives (*2*^*nd*^) (half siblings, *hs*; avuncular, *av*; grandparent-grandoffspring, *go*), 3^rd^ degree relatives (*3*^*rd*^) (first cousins, *fc*) and non-close kin. Every close kin category contains information about the number of possible effective dispersal events. For example, a pair of full siblings (*fs*) could have originated from the same breeding site from which each sibling moved into a gravitrap (two events), or they could have originated from different breeding sites (three events, including mother’s skip oviposition). Therefore, the corresponding number of possible dispersal events, for each case, can be calculated as the number of such breeding sites (n) plus one (n+1). Based on this, we constructed the sets with elements that represent the number of possible dispersal events for each case as {n_min_+1,.., n_max_+1}. For the *fs* category this set is FS = {2, 3}. In the case of a parent-offspring (*po*) pair, the minimum and maximum number of breeding sites is n_min_ = n_max_ = 1, giving a set PO = {2}. For the kinship category of 2^nd^ degree relatives, we have the following subsets: half siblings HS = {2, 3, 4, 5}; avuncular AV = {3, 4}; and grandparent-grandoffspring GO = {2, 3}. We constructed the full set for 2^nd^ degree relatives as the union of these three subsets (containing unique elements): 2ND = HS ⋃ AV ⋃ GO = {2, 3, 4, 5}. In the case of 3^rd^ degree relatives (first cousins *fc*), the minimum number of contributing breeding sites is n_min_ = 1 (first cousins and their mothers all originate from the same breeding site), while the maximum is n_max_ = 4 (first cousins and their mothers each originate from a unique breeding site). Therefore, for the category of 3^rd^ degree relatives we can construct a set as 3RD = FC = {2, 3, 4, 5}.

We then created a set of possible effective dispersal distances for each detected close kin pair based on their distance and assigned kinship category. This set of distances contains the same number of elements as the corresponding set of possible effective dispersal events (described above) and its values are obtained by dividing the detected spatial distance between a pair (d) with the corresponding set element. For example, if a collected pair AB, separated by distance d_AB_, falls into the category of 3^rd^ degree relatives, then a set of possible effective dispersal distances for this pair would be d_3rd_,_AB_ = {d_AB_/2, d_AB_/3, d_AB_/4, d_AB_/5}. For a full-sibling pair BC separated by spatial distance d_BC_, the set of possible effective dispersal distances will be d_fs,BC_={d_BC_/2, d_BC_/3}.

By combining the values from all pairwise sets of possible effective dispersal distances into one dataset, we characterized the resulting distribution of possible effective dispersal distances. This dataset was used to generate a probability density function (*pdf*) of effective dispersal distance (i.e. effective dispersal distance kernel) by fitting different functions (exponential, Weibull, log-normal) using the R package ‘fitdistrplus’ [41] that incorporates maximum likelihood estimation and parametric bootstrapping to generate median as well as 2.5 and 97.5 percentiles of each distribution parameter. To determine the ‘best fitting’ of the tested distributions, we assessed the Q-Q plots and computed goodness-of-fit statistics with an approximate Kolmogorov-Smirnov test, Akaike Information Criterion (AIC) and Bayesian Information Criterion (BIC) [41].

To estimate a *pdf* for randomly spaced individuals across the sampled area, we simulated 100 datasets where pairs had a randomly assigned kinship category and a random distance based on the available trap locations. The number of simulated pairs in each kinship category matched the number of observed pairs in the empirical dataset. We then applied the analytical procedure described above on all simulated datasets, and compared the simulated (random) and empirical distributions in the R package ‘sm’ [42] using the permutation test of equality of two distribution densities.

### Isolation-by-distance analysis and estimation of the dispersal kernel spread

When the probability of dispersal declines with distance, a positive correlation between genetic and geographic distances between individuals is expected, and this relationship is known as isolation-by-distance (IBD) [43]. IBD analysis can be used even when few or no close kin are captured. In fact, highly-related individuals should to be removed from the IBD analysis in order for it to reflect the long-term population processes [44], and we created a subsample for each area by removing individuals identified as close-kin, leaving 63 and 85 individuals in the Tampines and Yishun subsample, respectively. The significance of IBD was tested separately in Tampines and Yishun using Mantel’s correlation test with 1000 permutations, as implemented in the R package ‘ecodist’ [45].

IBD is best illustrated by the regression of pairwise genetic distances onto geographic distances among individuals [46]. The slope of this linear regression and the effective density can be used to estimate the standard deviation of the dispersal kernel (σ), that is also known as the dispersal kernel spread. The dispersal kernel spread can be calculated as: σ = √(1/4**π***Db*), where *b* is the slope of the linear regression and *D* is the effective density of reproducing individuals.

The slope of the linear relationship was estimated using the lm() function in R (R Core Team) for three different sets of genetic and geographic matrices. The first set included a matrix of PCA-based genetic distances against the matrix of linear geographic distances. A matrix of Rousset’s *â* or Loiselle’s kinship coefficient was tested against the matrix of ln-transformed geographic distances, given that both genetic estimators are expected to vary approximately linearly with the natural logarithm of the distance [38].

The effective density *D* is defined as *N*_*e*_/study area, where *N*_*e*_ is the effective population size. We estimated *N*_*e*_ using two genetic methods based on a single sample. The first method is *N*_*e*_ estimation by Waples and Waples [47], based on the ‘parentage analysis without parents’ (method 1, PWoP) that uses the frequency of full-siblings and half-sibs in a population sample to reconstruct the number of parents that contributed to such a sample. The median and the 95% confidence interval were calculated using 100 resamples with a random replacement of one individual. The second method represents *N*_*e*_ estimation by Waples and England [48] that is based on the linkage disequilibrium data (method 2, LD_Ne_), with the 95% confidence interval calculated using the jackknifing procedure over loci implemented in the program *N*_*e*_ estimator v.2.1 [49]. Finally, effective density was estimated using the entomological survey data from the Gravitrap sentinel trap system across the study areas (method 3, Gravitrap). The numbers of adult females caught from January 2018 through May 2018 were used to estimate the average number of breeding females per square meter and this number was multiplied by 2 to give the effective number of breeding individuals per unit area, as we assume 1:1 sex ratio in an *Ae. aegypti* population [50]. For Tampines, we considered patch 1 (larger sampling area) as a more representative population sample for the calculations of local *N*_*e*_ and density than the smaller patch 2 (Suppl. Fig.1A).

### Spatial autocorrelation analysis

Under spatially limited dispersal and breeding, the population is expected to develop a patchy distribution of genotypes, with positive spatial autocorrelation declining with distance [51,52]. The point at which the correlogram curve crosses the x-axis provides an estimate of the genetic patch size [53]. To compute the correlogram curve for each sampling site, we used PCA genetic distance among all genotyped mosquitoes in a site, and the spatial grouping within distance classes that were incrementally increased by 50 m. The analysis was done in GenAlEx v.6.501 [54]. The autocorrelation coefficient under the null hypothesis of no spatial structure was generated by the permutation procedure that shuffles all individuals among the geographic locations within a site 1000 times and generates 95% CI with the 25^th^ and 975^th^ ranked permutated values. 95% CI for the observed autocorrelation coefficient for each distance class was obtained from 1000 random draws of individuals with replacement [55].

## Results

### Spatial distribution of close-kin pairs

In the total dataset that included 107 individuals from Tampines and 108 from Yishun, we detected 76 close-kin (CK) pairs: 19 full-sibling pairs, 18 pairs of 2^nd^ degree relatives, and 39 pairs of 3^rd^ degree relatives (Supplementary Table 1). We did not detect any parent-offspring (*po*) pairs, and 26.3% (20/76) of CK pairs were found within a building (Figure 1A) – on the same floor or 4-5 floors apart (19/20 pairs). Nearly half (47%) of all detected full-sibling pairs were caught within a building, in comparison to 27.8% of all 2^nd^ degree relatives and 15.4% of all 3^rd^ degree relatives (Figure 1B).

**Figure 1.**
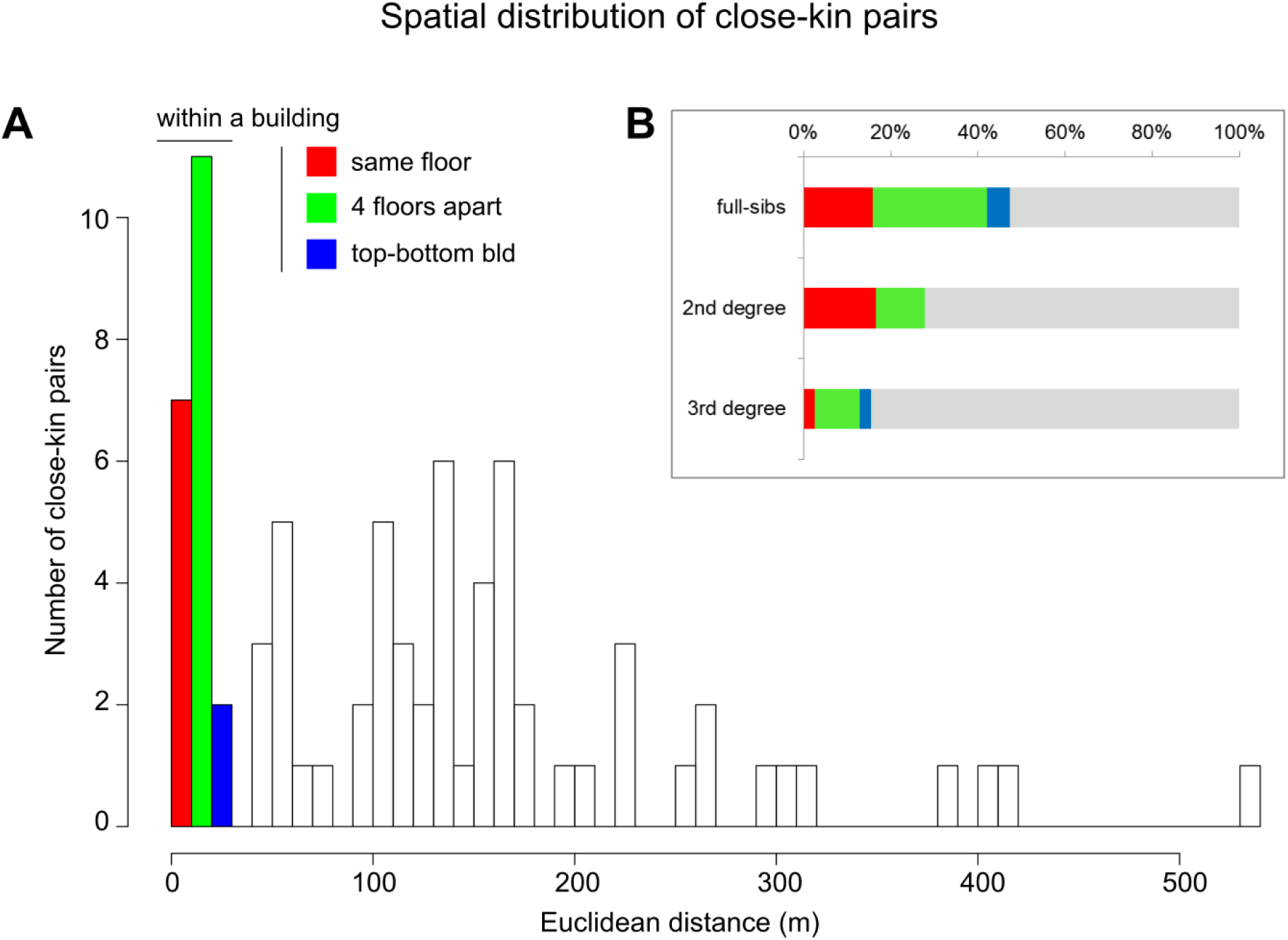
Spatial distribution of close-kin (CK) pairs. **A.** Histogram of the number of CK pairs detected in Tampines and Yishun, with their pairwise separation distances binned into the 10 m categories. The first three bins include pairwise distances within a building (where CK are found on the same floor, or 4-5 floors apart, or at the top and bottom of a building). **B.** Percentage of the total number of pairs in each kinship category (full-sibling, 2^nd^ and 3^rd^ degree relatives) found within the same building or in different buildings.

The mean pairwise distance for CK was 124.3 m (full-sibling = 107.3 m, 2^nd^ degree = 135 m, 3^rd^ degree = 127.4 m), with 90% of CK found up to 264 m apart, and the largest distance of 531 m (for one full-sibling pair), which points to a long-tailed dispersal kernel (with rare long-range dispersal events).

### Dispersal kernel parametrization from close-kin data

The ‘goodness of the fit’ criteria (AIC, BIC, the Kolmogorov-Smirnov statistic, Table 1) and the Q-Q plot (Suppl. Fig. 2) indicated that the distribution of possible effective dispersal distances is well described by the Weibull and negative exponential (Laplacian) distribution. Given that the estimated Weibull shape parameter (*k*) was close to 1 (median 1.11, 95% CI: 1.01-1.22), the Weibull distribution can be reduced to an exponential distribution with the estimated rate parameter λ = 0.022 (95% CI: 0.020-0.025) (Table 1). This rate parameter for the combined dataset (all CK) was highly congruent with the estimates from separate CK categories: full-sibslings λ = 0.019 (95% CI: 0.014-0.027), 2^nd^ degree relatives λ= 0.021 (95%CI: 0.016-0.027), 3^rd^ degree relatives λ= 0.024 (95%CI: 0.020-0.028) (Figure 2). Both the mean and standard deviation (σ) of the exponentially distributed effective dispersal distance are 1/λ = 45.2 m (95%CI: 39.7-51.3 m), and the inferred dispersal kernel gives the 95% probability of effective dispersal distance up to 136 m (95%CI: 120-152 m).

**Table 1.** Goodness-of-fit statistic and criteria, and parameter estimates from the distribution fitting analysis for the close-kin data. The Kolmogorov-Smirnov statistic measures the distance between the fitted parametric distribution and the empirical distribution, and AIC and BIC assess the model fit while penalizing the number of estimated parameters. A lower value of the statistic or either criterion indicates a better fit. Median and 95% CI for the parameters were generated with 1000 bootstraps.

| Goodness-of-fit test | exponential<br>(Laplacian) | Weibull | log-normal |
| --- | --- | --- | --- |
| Kolmogorov-Smirnov | 0.084 | 0.057 | 0.125 |
| AIC | 2387.913 | 2386.055 | 2429.829 |
| BIC | 2391.427 | 2393.082 | 2436.856 |
| Parameter | $\lambda$ | shape | meanlog |
| median (95% CI) | 0.022 (0.020-0.025) | 1.108 (1.011-1.222) | 3.322 (3.170-3.462) |
|  | - | scale<br>46.939 (41.770-52.445) | sdlog<br>1.162 (1.055-1.252) |

**Figure 2.**
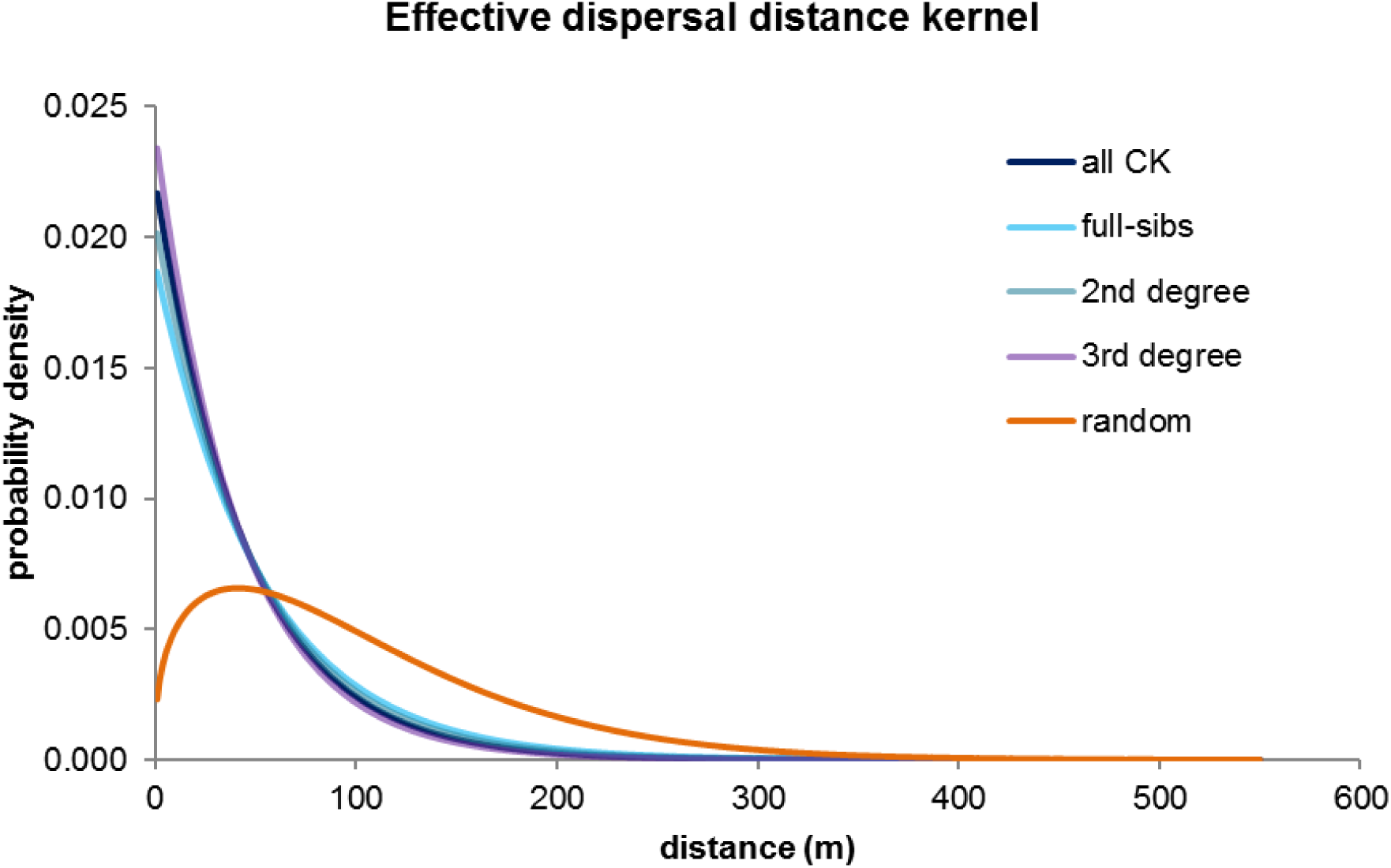
Effective dispersal distance kernel estimated from the close-kin data. The inferred *pdf*s are highly congruent among separate datasets (full-sibling, 2^nd^ and 3^rd^ degree relatives) and the combined dataset (all CK), and are significantly different from the simulated dataset (random) that reflects random sampling from the available traps.

Estimated probability density function (*pdf*) for the random distribution of pairwise close-kin distances in this landscape had significantly different parameters (Weibull shape *k* = 1.351, 95% CI: 1.338-1.364; Weibull scale λ = 111.630, 95%CI: 110.502-112.696) from the empirically observed data (permutation test for equality of densities *p* < 0.001), indicating that the simulated (random) dataset reflects the spatial distribution of traps rather than intentional effective dispersal (Figure 2).

### Dispersal kernel spread from the IBD analysis

The three genetic distance measures (PCA, Rousset *â*, Loiselle’s kinship) gave highly congruent results in the IBD analysis (Suppl. Table 1), and we focus on the results with the PCA genetic distance in the main text.

Significant IBD was detected in both Tampines (Mantel *r* = 0.124, 95%CI: 0.052-0.198; *R*^*2*^ = 0.015, *p* = 3.91*10^−8^) and Yishun (Mantel *r* = 0.158, 95%CI: 0.112-0.208; *R*^*2*^ = 0.024, *p* = 2.18*10^−21^) (Figure 3). The estimated IBD slope *b* was 0.0037 (95% CI: 0.0024-0.0050) for the Tampines data, and 0.0065 (95% CI: 0.0051-0.0079) for the Yishun dataset (Table 2). The estimated mean effective population density *D* varied from 0.0014 to 0.0074 for Tampines, and from 0.0014 to 0.0063 for Yishun, depending on the method for effective population size estimation (methods 1 and 2), or the entomological survey data (method 3) (Table 2).

**Table 2.** IBD-based estimates of the dispersal kernel spread (σ). The mean (95% CI) for IBD slope (*b*), effective population size (*N*_*e*_), effective density (*D*) estimated using the methods 1-3, and the dispersal kernel spread (σ) for *Aedes aegypti* data from Tampines and Yishun.

| Tampines |  |  |  |  |
| --- | --- | --- | --- | --- |
| $b$ | | $N_e$ | $D$ | $\sigma$ |
| 0.0037 (0.0024-0.0050) | method 1<br>(PWoP) | 863 (863-1112) | 0.0074 (0.0074-0.0095) | 54.1 m (40.8-67.7 m) |
| | method 2<br>(LD$N_e$) | 167 (93-619) | 0.0014 (0.0007-0.0053) | 122.9 m (54.7-206.2 m) |
|  | method 3<br>(Gravitrapp) | - | 0.0048 (0.0030-0.0066) | 66.8 m (48.8-105.6 m) |
| Yishun |  |  |  |  |
| $b$ | | $N_e$ | $D$ | $\sigma$ |
| 0.0065 (0.0052-0.0079) | method 1<br>(PWoP) | 1185 (1176-1346) | 0.0063 (0.0063-0.0072) | 44 m (37.6-49.7 m) |
| | method 2<br>(LD$N_e$) | 258 (200-357) | 0.0014 (0.0011-0.0019) | 94.4 m (73-120.6 m) |
|  | method 3<br>(Gravitrapp) | - | 0.0022 (0.0015-0.0030) | 74.4 m (57.9-104.9 m) |

**Figure 3.**
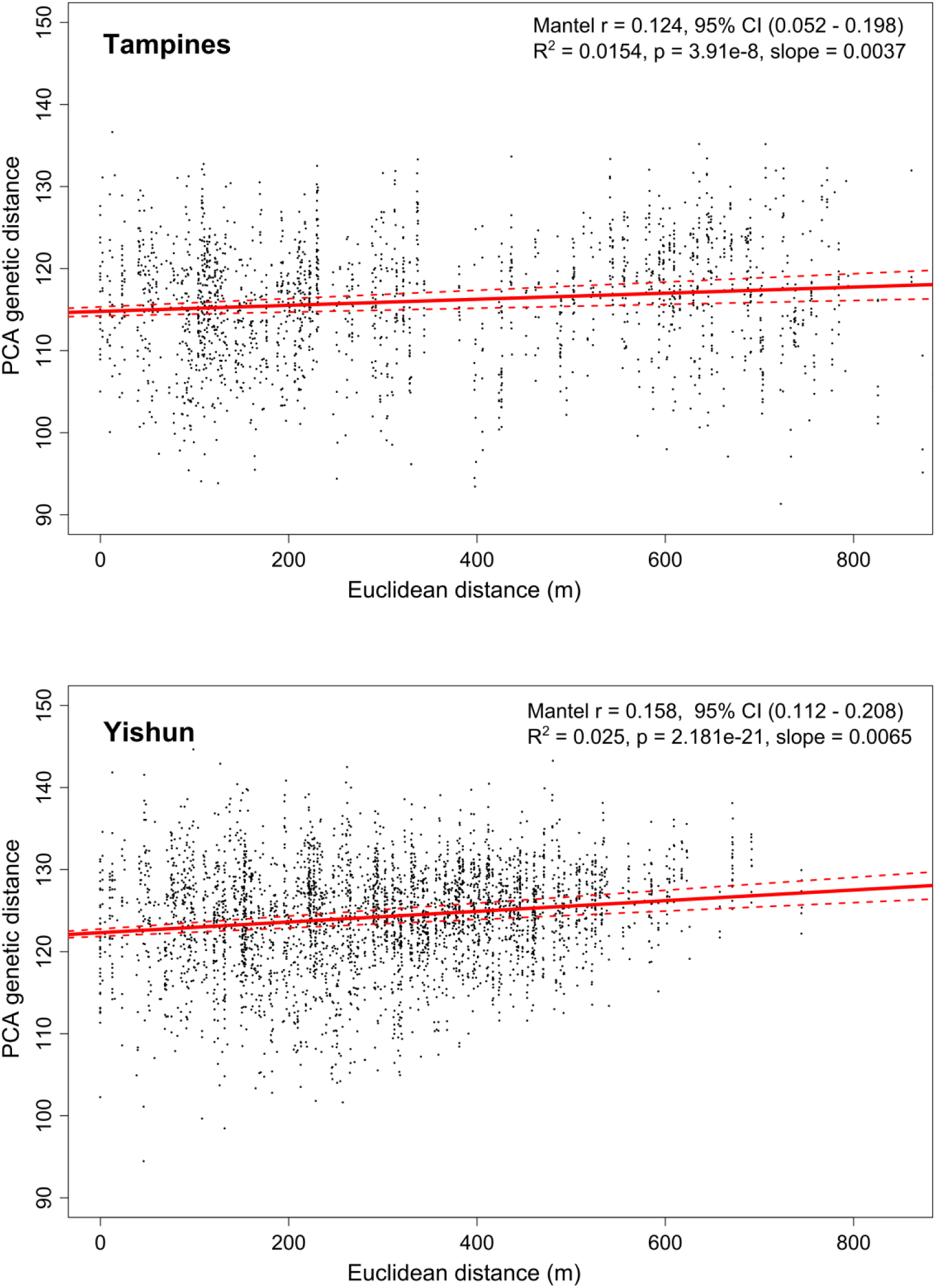
Isolation-by-distance analysis on datasets from Tampines (upper) and Yishun (lower). Mantel test and linear regression were applied to the matrices of PCA genetic distance and linear geographic distance between pairs of individuals, with close kin removed from both datasets. The red line shows regression with 95% CI (dashed line).

Taking into account the uncertainty of both parameter estimates (95% CI for *b* and *D*), the estimated effective dispersal kernel spread (σ) for Tampines was 54.1 m (40.8-67.7 m, method 1 PWoP), 122.3 m (54.7-206.2 m, method 2 LD_Ne_), 66.8 m (48.8-105.6 m, method 3 gravitrap). For Yishun, the estimated σ was 44 m (37.6-49.7 m, method 1), 94 m (73-120.6 m, method 2), 74.4 m (57.9-104.9 m, method 3) (Table 2, Figure 4). It is worth noting a good overlap between the effective density estimates from the genetic data and the entomological data from the gravitraps (method 3) that preferentially target the ovipositing females (Figure 4).

**Figure 4.**
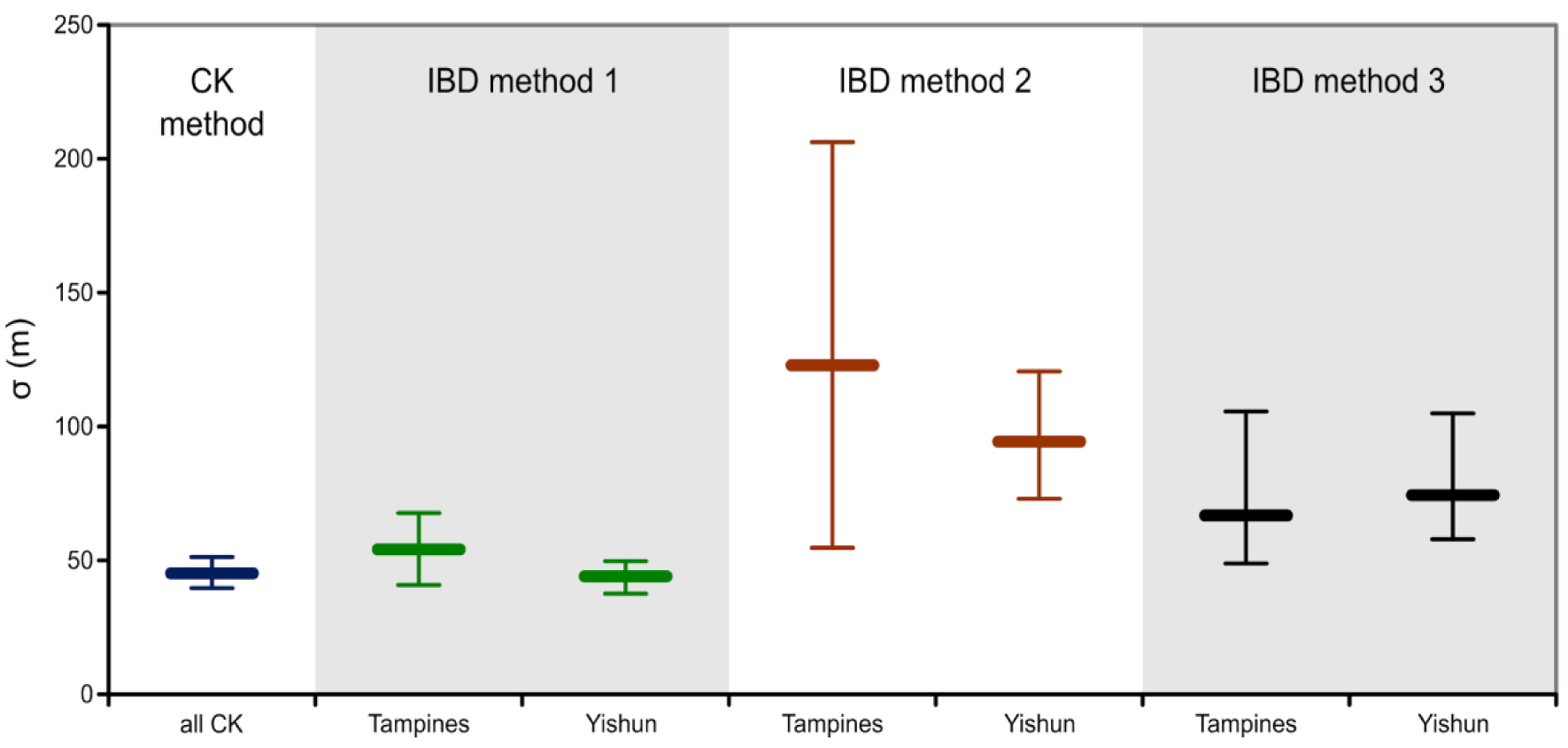
The dispersal kernel spread (σ) estimated from the close-kin data and IBD analysis. Sigma (σ) and its 95% CI are plotted for the combined close-kin data (CK method) and PCA-based IBD analysis for Tampines and Yishun (with effective density estimates from methods 1-3).

Applying the exponential dispersal kernel (found to fit the close-kin data), where σ represents both standard deviation and the mean, the estimates of σ from the IBD analysis are equivalent to the mean effective dispersal distance and can be used to parametrize the *pdf* with 1/σ = λ, assuming isotropic dispersal in two dimensions (Broquet and Petit 2009). Our results indicate that the dispersal kernel parameter (σ) estimated from the close-kin data and indirectly through the IBD analysis can yield similar results (Table 2, Figure 4).

### Spatial autocorrelation analysis - genetic patch size

Significantly positive spatial autocorrelation at distances up to 200 m, with the highest correlation coefficient within the first 50 m, was detected in both Tampines and Yishun (Figure 5). This indicates that individuals found up to 200 m from each other are more genetically similar than if randomly sampled across 750 m, with the highest genetic similarity (relatedness) within 50 m from each other. Significantly negative autocorrelation was detected at 300 m in Yishun and 500 m in Tampines, putting the x-intercept between 200 m and 300 m in Yishun and between 200 m and 500 m in Tampines. The x-intercept for a squared sampling area closely approximates the length of one side of a ‘genetic patch’ [53], delineating a ‘genetic patch’ area of at least 200 × 200 m in this landscape. The spatial extent of the genetic patch also reflects the spatial extent of the effective dispersal distance kernel ‘tail’; e.g. 99^th^ percentile of the *pdf* (derived from the close-kin data) was at 206 m (95%CI: 184-234 m).

**Figure 5.**
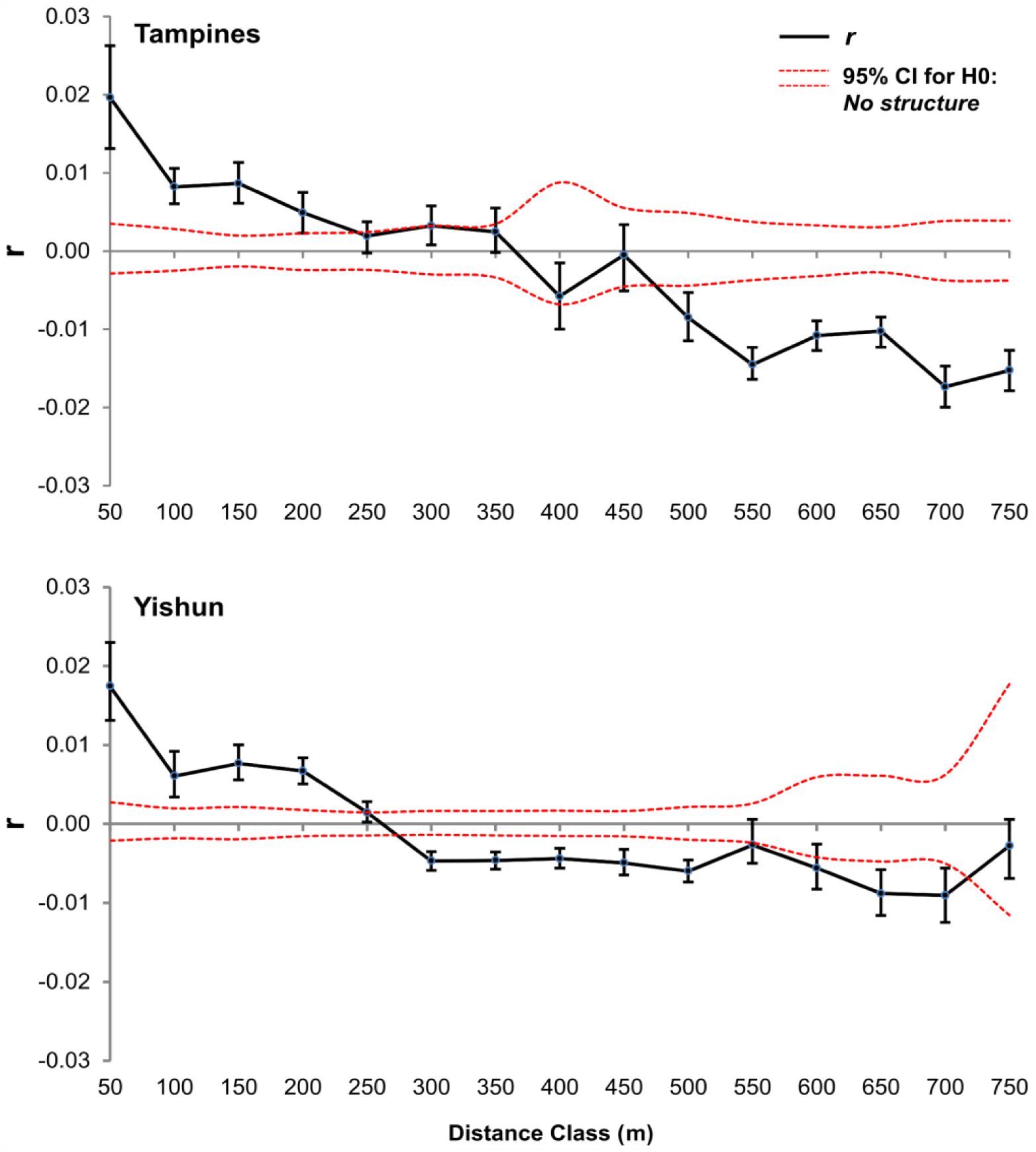
Spatial genetic autocorrelation in Tampines and Yishun. The ending point of a distance class is on the x-axis, and spatial autocorrelation coefficient (r) of genotypes in Tampines (107 individuals) and Yishun (108 individuals) is on the y-axis. Two dashed lines along the x-axis are the permutated 95% CI of autocorrelations under the null hypothesis of a random distribution of genotypes in space. Vertical lines are the bootstrapped 95% CIs with mean genetic autocorrelation.

## Discussion

Here we show how the analyses of spatial genetic patterns can be used to characterize the effective dispersal of a mosquito like *Ae. aegypti*, and we discuss the utility of this approach in an operational context.

Our newly developed method allows for the parametrization of the effective dispersal distance kernel, as it decomposes the observed distances between close-kin to generate the distribution of potential effective dispersal distances (achieved over one generation of successful reproduction). It gives probabilities of dispersal distances in any direction, that Nathan et al. (2012) [56] refer to as ‘dispersal distance kernel’ (k_D_(r)), and it should not be confused with the ‘dispersal location kernel’ (k_L_(r)) that gives probabilities for the end locations of dispersers relative to the source locations [56]. Location kernel can be derived from the distance kernel and *vice versa*, given their relation: k_D_(r) = 2**π**rk_L_(r) in a two-dimensional habitat [56], and k_D_(r) = 4**π**r^2^k_L_(r) in a three-dimensional habitat.

The exponential (Laplacian) kernel inferred from the close-kin data in Yishun and Tampines gives 50% probability of an effective dispersal occurring within 32 m, which indicates that, in this landscape we can expect half of the successful breeders to stay within the high-rise building where they emerged or move to the adjacent building. The mean effective dispersal distance (and dispersal kernel spread σ) was estimated at 45.2 m (95%CI: 39.7-51.3 m), with a 10% probability of a dispersal distance greater than 100 m (95%CI: 92-117 m). Our genetic-based estimates match the parametrized dispersal kernel from mark-release-recapture (MRR) experiments performed in Brazil and Malaysia with *Ae. aegypti* males from a genetically engineered line OX513A [7]. Namely, their dispersal kernel gave estimates of a high level of dispersal to up 33 m, a mean distance travelled of 52.8 m (95% CI: 49.9-56.8 m) in Brazil and 58 m (95%CI: 52.1-71 m) in Malaysia, with 10% of dispersers moving >105.7 m (95%CI: 86.5-141.1 m) [7]. Moreover, globally collated MRR experimental data for *Ae. aegypti* [14] produced an exponential kernel with σ = 54.1 m [10]. Such high congruence with our genetic-based inferences indicates the robustness of spatial-genetic patterns in reflecting the dispersal characteristics of this mosquito, and it demonstrates the utility of our genetic-based method as a viable alternative to conventional, operationally-demanding MRR experiments.

Using the IBD approach, we obtained a wider range of dispersal spread values (σ) when compared to our close-kin approach, depending on which method was used to estimate the effective population size (*N*_*e*_) and density. Specifically, the highest congruence between the IBD and the close-kin analyses was obtained using the *N*_*e*_ estimates from the PWoP method (method 1, Table 2), that gave σ of 54.1 m (95% CI: 40.8-67.7 m) for Tampines and 44 m (95% CI: 37.6-49.7 m) for Yishun. The second best match was obtained using the entomological effective density estimate (the number of gravitrap-caught mosquitoes/m^2^, method 3), that gave σ of 66.8 m (95% CI: 48.8-105.6 m) for Tampines and 74.4 m (95% CI:57.9-104.9 m) for Yishun. The linkage disequilibrium-based *N*_*e*_ estimate (method 2) produced the widest range of σ values (Tampines 54.7-206.2 m, Yishun 73-120.6 m) (Figure 4). Two conclusions can be drawn from these results. First, the IBD measure of the dispersal kernel spread represents a long-term average but it can match the short-term measure from the close-kin analysis. Given that close-kin method requires more intensive sampling in order to capture enough close-kin pairs for the reliable kernel parametrization, an opportunity to use the IBD approach instead is appealing, particularly under budgetary limitations for genome-wide genotyping. However, IBD-based estimation of σ requires accurate estimation of effective population size, which it is not easily achievable [57] and the uncertainty about this parameter has more impact than the uncertainty in the IBD slope [58]. Moreover, the IBD method assumes long-term stability of mosquito dispersal patterns and abundance and is therefore a meaningful alternative to the close-kin method in populations that do not experience strong seasonal fluctuations, landscape alterations, intensive control campaigns etc. Second, our analyses suggest that the gravitrap data reflects the breeding rather than the census population size, and could be used as an entomological proxy for the effective population size, complementing the genetic based estimation of this population parameter.

In Tampines and Yishun, that are largely homogeneous landscapes with multi-storey apartment buildings, 47% of all detected full-siblings were found on the same floor or 4-5 floors apart, and the inferred dispersal kernel gives the expectation of high level of effective dispersal within a building or between adjacent buildings. This is in agreement with a previous study in Singapore where *Ae. aegypti* females marked with rubidium *via* blood meal were released from middle floors and moved readily towards the top or bottom of multi-storey buildings in search of oviposition sites [59]. In Trinidad, significantly more eggs were collected in ovitraps 13–24 m above ground level than at any other elevation [60], again suggesting that vertical movement is common.

Given that the high-rise apartment blocks can provide an abundance of hosts, oviposition and resting sites for *Ae. aegypti*, their tendency to remain close to the birthplace is not unexpected. The tail of the dispersal kernel, however, provides insight into rarer long-range dispersal events that are consequential for the control strategies that rely on the releases of modified mosquitoes for population suppression or replacement. For example, the speed of the *Wolbachia* infection spread is expected to be slower in *Ae. aegypti* populations with longer-tail dispersal kernels, but this dispersal pattern also allows the initiation of the *Wolbachia* spread with smaller local releases [61]. Based on our parametrized dispersal kernel and theoretical approximation [61], the spread of *Wolbachia* (with a fitness cost equivalent to *w*Mel strain) in a landscape like Tampines and Yishun could be initiated with the releases of *Ae. aegypti* in a circular area with a radius of at least 100-130 m (2.51σ). Diameter of this area (200-260 m) also corresponds to the length of one side of a ‘genetic patch’ estimated with the spatial autocorrelation analysis (∼200-300 m) that reflects the extent of the effective dispersal kernel’s ‘tail’ (e.g. 99^th^ percentile = 206 m, 95%CI: 184-234 m).

The above mentioned theoretical approximation of the conditions for *Wolbachia* initiation and spread [61] assumes isotropic dispersal in a 2-dimensional habitat. Optimal release strategy for the Sterile Insect Technique programs [62], different suppression and replacement strategies [63,64], the effect of larval habitat fragmentation on population crash [65], and confinement and reversibility conditions for threshold-dependent gene drive systems [10] have all been simulated in the spatially explicit models of mosquito populations that applied the exponential dispersal kernel in a 2-dimensional landscape. The approximation of the parametrized dispersal kernel for high-rise landscapes could be achieved by considering the releases of mosquitoes from multiple floors rather than from the ground level only. Clearly, further theoretical and simulation modelling development that incorporates mosquito dispersal characteristics in a 3-dimensional, heterogeneous habitat is needed to robustly predict the requirements and outcomes of different mosquito control strategies in landscapes with a prominent vertical dimension.

## Conclusion

Accurate characterization of dispersal patterns in the field is critical for the optimization of surveillance and control of disease vectors like *Ae. aegypti.* Knowledge of the dispersal kernel parameters enables operational teams to design and implement optimal coverage of monitoring or release points in a given landscape, facilitating the efficient utilization of resources and maximizing the impacts of interventions. We show that spatial genetic analyses can provide robust estimates of mosquito dispersal patterns, with our newly developed method for the construction of the effective dispersal distance kernel through close-kin analysis enabling the most comprehensive estimate of relevant parameters. The indirect inference from the isolation-by-distance framework, that requires less intensive sampling than close-kin analysis, can provide estimates of dispersal kernel spread; however, this approach is sensitive to the inaccurate estimates of effective population size and is uninformative about the probabilities of long-range dispersal that can have important implications for control programs. IBD analysis can be complemented with spatial autocorrelation analysis to ascertain the spatial extent of the effective dispersal kernel ‘tail’ through estimation of the genetic patch size. With the decreasing cost of next generation sequencing, acquisition of spatial genetic data will become increasingly accessible, and given the complexities and criticisms of conventional MRR methods, but the central role of dispersal measures in essential vector control programs, we recommend genetic-based dispersal characterization as the more desirable means of parameterization.

## Acknowledgments

We would like to thank Dr. Chee Seng Chong and the colleagues from the Applied Entomology and Informatics Sections at EHI, for providing mosquito specimens and entomological data, and Mr. Chew Ming Fai, Director-General for Public Health, National Environment Agency (NEA) and Associate Prof. Lee Ching Ng, Director, EHI for their support and approval to publish the study. We also thank Dr. Paola Florez de Sessions, Dr. Yuan Zhu, Dr. Chiea Chuen Khor, Dr. Chih Chuan Shih and colleagues at Genome Institute Singapore, A* Star, Singapore for their assistance in sequencing the RADseq libraries.

## Author Contributions

Conceptualization: RG, HHC, FI. Methodology: RG, FI. Formal Analysis: FI, RG. Field Work and Data Acquisition: TWP, MABAR, CL, TCH, HHC. Data Curation: HHC, FI. Project Administration: HHC. Writing – Original Draft Preparation: RG, FI. Writing – Review and Editing: HHC, DGJ, RG. Funding Acquisition: HHC, DGJ, RG.

## Supplemental Figures and Tables

**Supplemental Figure 1.**
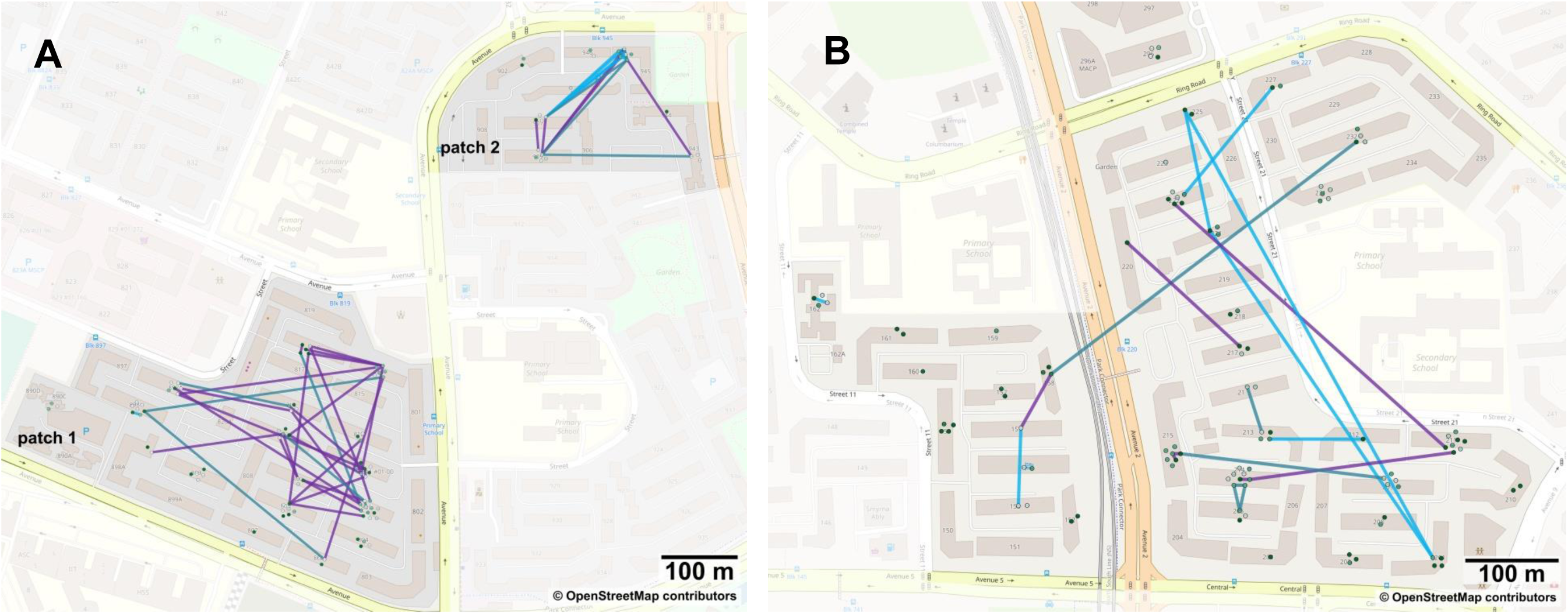
Spatial network of close kin in Tampines (A) and Yishun (B). Cyan lines connect full siblings, blue-green lines connect 2^nd^ degree relatives, and purple lines connect 3^rd^ degree relatives. Gravitraps are represented with green dots - one group per building, with shades varying from lighter to darker depending on the altitude (light green - ground floor, dark green - top floor).

**Supplemental Figure 2.**
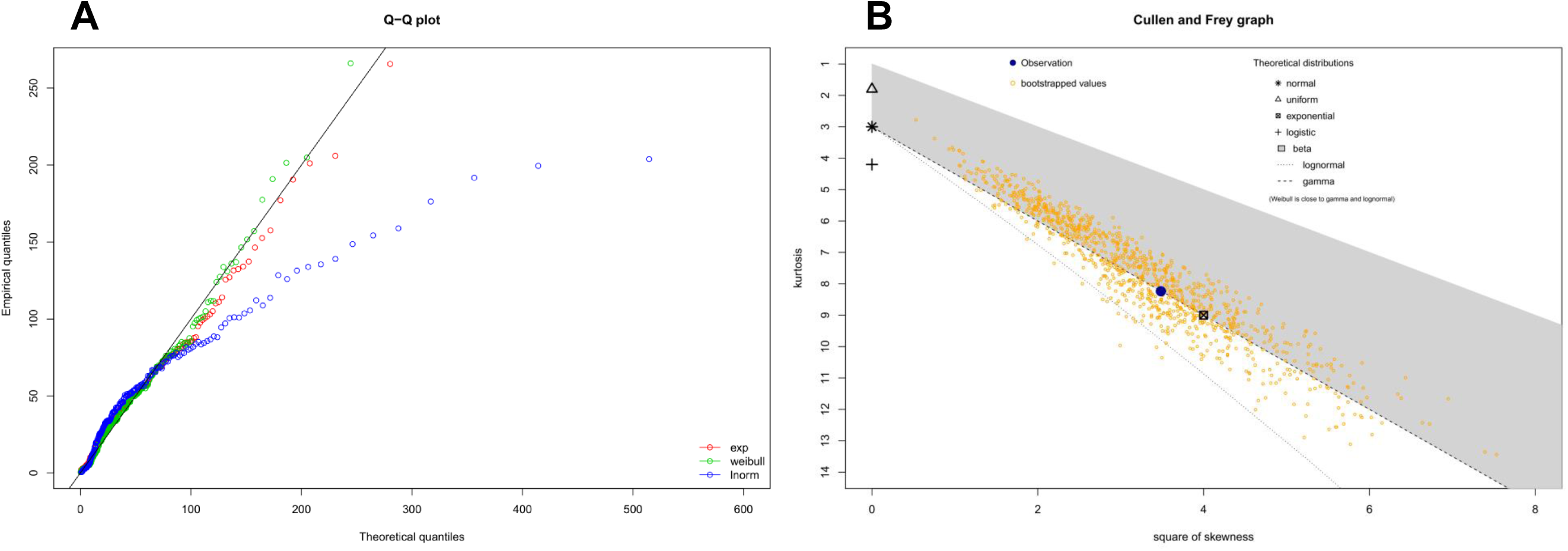
Distribution fitting (dispersal kernel parametrization) analysis in the R package fitdistrplus [1]. **(A)** Q-Q plot for the empirical and theoretical distribution quantiles This goodness-of-the-fit assessment shows a good fit of the empirical to the expected data for Weibull (green) and exponential (red) distributions, but not to the lognormal distribution (blue). **(B)** Skewness-kurtosis plot. A nonparametric bootstrap procedure (constructed by random sampling with replacement from the empirical data set) was performed to take into account the uncertainty of the estimated values of kurtosis and skewness from data.

---

**Supplemental Table 1.**
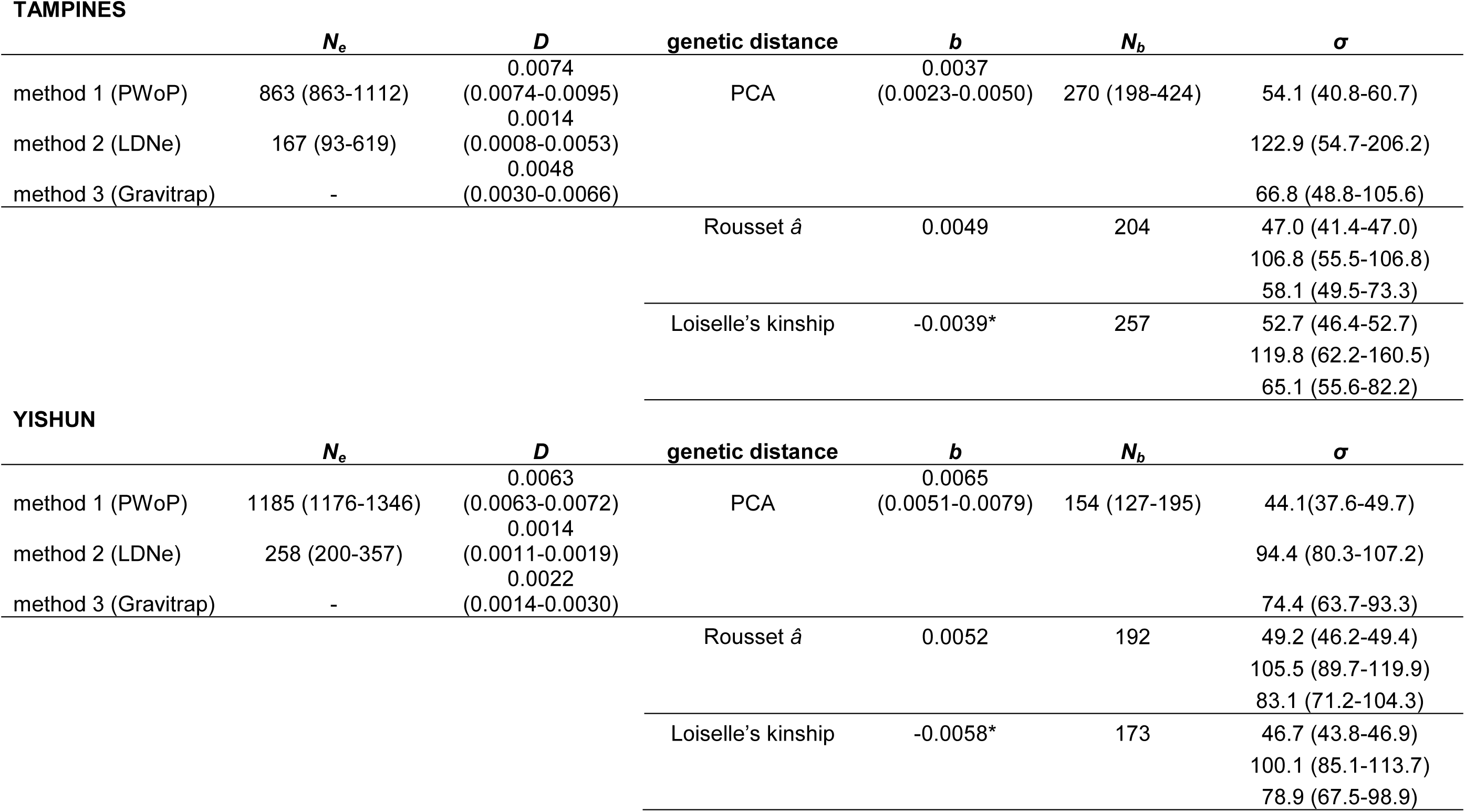
IBD-based estimates of the dispersal kernel spread (σ) in *Aedes aegypti* from Singapore. The mean (95% CI) for estimates of: effective population size (*N*_*e*_) and effective density (*D*) calculated using 3 methods (genetic – PWoP, LDNe; entomological – Gravitrap), and IBD slope (*b)* and genetic neighborhood size (*N*_*b*_) calculated from the analysis with different genetic distance measures (PCA, Rousset *â*, Loiselle’s kinship), for data from Tampines and Yishun. *Note the negative relationship between Loiselle’s kinship coefficient and ln-geographic distance (i.e. kinship decreases as the ln-geo distance increases).

**TAMPINES**
| | $N_e$ | $D$ | genetic distance | $b$ | $N_b$ | $\sigma$ |
| --- | --- | --- | --- | --- | --- | --- |
| method 1 (PWO) | 863 (863-1112) | 0.0074<br>(0.0074-0.0095) | PCA | 0.0037<br>(0.0023-0.0050) | 270 (198-424) | 54.1 (40.8-60.7) |
| method 2 (LDNe) | 167 (93-619) | 0.0014<br>(0.0008-0.0053) |  |  |  | 122.9 (54.7-206.2) |
| method 3 (Gravitrapp) | - | 0.0048<br>(0.0030-0.0066) |  |  |  | 66.8 (48.8-105.6) |
| | | | Rousset $\hat{a}$ | 0.0049 | 204 | 47.0 (41.4-47.0) |
|  |  |  |  |  |  | 106.8 (55.5-106.8) |
|  |  |  |  |  |  | 58.1 (49.5-73.3) |
|  |  |  | Loiselle’s kinship | -0.0039* | 257 | 52.7 (46.4-52.7) |
|  |  |  |  |  |  | 119.8 (62.2-160.5) |
|  |  |  |  |  |  | 65.1 (55.6-82.2) |

**YISHUN**
| | $N_e$ | $D$ | genetic distance | $b$ | $N_b$ | $\sigma$ |
| --- | --- | --- | --- | --- | --- | --- |
| method 1 (PWO) | 1185 (1176-1346) | 0.0063<br>(0.0063-0.0072) | PCA | 0.0065<br>(0.0051-0.0079) | 154 (127-195) | 44.1(37.6-49.7) |
| method 2 (LDNe) | 258 (200-357) | 0.0014<br>(0.0011-0.0019) |  |  |  | 94.4 (80.3-107.2) |
| method 3 (Gravitrapp) | - | 0.0022<br>(0.0014-0.0030) |  |  |  | 74.4 (63.7-93.3) |
| | | | Rousset $\hat{a}$ | 0.0052 | 192 | 49.2 (46.2-49.4) |
|  |  |  |  |  |  | 105.5 (89.7-119.9) |
|  |  |  |  |  |  | 83.1 (71.2-104.3) |
|  |  |  | Loiselle’s kinship | -0.0058* | 173 | 46.7 (43.8-46.9) |
|  |  |  |  |  |  | 100.1 (85.1-113.7) |
|  |  |  |  |  |  | 78.9 (67.5-98.9) |

